# Chromosome-level reference genome of the beach false foxglove, *Agalinis fasciculata* (Orobanchaceae)

**DOI:** 10.64898/2026.02.04.703813

**Authors:** Pedro H. Pezzi, Maribeth Latvis

## Abstract

Orobanchaceae is the largest family of parasitic plants, encompassing a full spectrum of parasitic strategies, ranging from autotrophic to holoparasitic. *Agalinis* is a genus of facultative hemiparasites comprising about 70 species distributed throughout the Americas, including several endemic and rare taxa. *Agalinis fasciculata*, the beach false foxglove, is a widely distributed species across southeastern North America. Here, we use PacBio HiFi, Omni-C, and RNA-seq data to generate the first high-quality reference genome for the genus. The nuclear genome is 2.29 Gb in size, with most sequences anchored to 14 pseudochromosomes and an N50 of 162 Mb. BUSCO analyses indicate high completeness (98.4%). Structural genome annotation identified 34,133 protein-coding genes and 39,266 transcripts, most of which have at least one functional annotation. The plastid and mitochondrial genomes were also assembled. We further examined genetic diversity and demographic history in *A. fasciculata*, revealing low genome-wide heterozygosity and evidence of inbreeding. This reference genome is an important resource for understanding the evolutionary history of the genus and the evolutionary patterns of parasitism within Orobanchaceae.

**Significance:** This high-quality genome is the first chromosome-level assembly for *Agalinis*, a hemiparasitic genus in the plant family Orobanchaceae. It improves the taxon sampling within Orobanchaceae, representing an important resource for investigating patterns of genome evolution in parasitic lineages. Furthermore, *Agalinis* has served as a focal genus for studies of the anatomy of haustorial development, and genome annotation incorporated RNA from multiple tissues, enabling the identification of genes expressed in different tissues, including roots and haustoria. This genome also serves as a reference for evolutionary studies of other *Agalinis* species, many of which are endemic and of conservation concern in North and South America. Overall, the beach false foxglove genome will support studies of the evolutionary history of *Agalinis* and genome evolution across Orobanchaceae.

## Introduction

Orobanchaceae is the largest family of parasitic plants, comprising over 2,000 species across 90 genera, with a wide range of reliance on host plants and photosynthetic capability, including autotrophy, hemiparasitism (photosynthetic), and holoparasitism (non-photosynthetic). With relaxed selective pressure on photosynthesis, increased reliance on host plants is accompanied by a dramatic reduction in vegetative structures, as well as parallel patterns of gene family loss and expansion (Xu et al. 2022). Currently, fewer than 20 species have publicly available genomes, representing 11 genera. Many Orobanchaceae are economically important as agricultural pests due to their parasitic lifestyle, and about half of the available genomes come from pest genera. However, genomic resources remain limited for species that do not parasitize crops. An improved taxon sampling is crucial for understanding genome evolution across the evolutionary breadth of the family.

Within Orobanchaceae, *Agalinis* (the false foxgloves) is a hemiparasitic genus distributed across the Americas, likely originating in North America and later dispersing into South America, accompanied by increased diversification and pollinator and life-history strategy shifts (Pezzi et al. 2026). Of the ∼70 described species, which now includes the genus *Esterhazya* (Latvis et al. 2024; Souza et al. 2025), many have highly restricted distribution, and in North America, several species are considered imperiled in at least one state. Moreover, *Agalinis* served as a focal genus in early anatomical studies on the induction of haustoria, the specialized roots of plant parasites that penetrate host plants, with implications for host recognition (Chang and Lynn 1986). A reference genome for *Agalinis* represents a valuable resource for conservation genomic studies of imperiled species in North and South America, as well as for understanding genome evolution and selective pressures in hemiparasitic taxa.

We present a chromosome-level genome of the beach false foxglove, *A. fasciculata* (Elliott) Raf.. We assembled the nuclear genome and annotated it using RNA-seq data from flowers, roots, and leaves. We also estimated the genetic diversity and reconstructed the historical demographic history of the species. The addition of an Orobanchaceae reference genome will advance phylogenomic comparative analyses of parasitism and provide a valuable resource for studies of *Agalinis* evolution and conservation.

## Results and Discussion

### Sequencing and Nuclear Genome Assembly and Scaffolding

The newly assembled nuclear genome of *A. fasciculata* has a total length of 2.29 Gb, with an N50 of 162 Mb and an N90 of 131 Mb (Fig. 1a). The genome was estimated to be ∼2.3 Gb (Fig. S1), diploid (Fig. S2), and showed low heterozygosity (0.15%) and high repetitive content (80.4%). After decontamination (Fig. 1b), the assembly comprised 153 scaffolds. Of these, the 14 longest scaffolds corresponded to the 14 chromosomes, which agrees with previous estimates for the base chromosome number of this species (Canne 1984). We identified at least one telomeric repeat on each pseudochromosome (Fig. S3). Together, these 14 pseudochromosomes encompass 98.89% of the total genome length. The Omni-C contact map shows clear chromosome-scale scaffolds (Fig. S4).

**Figure 1.**
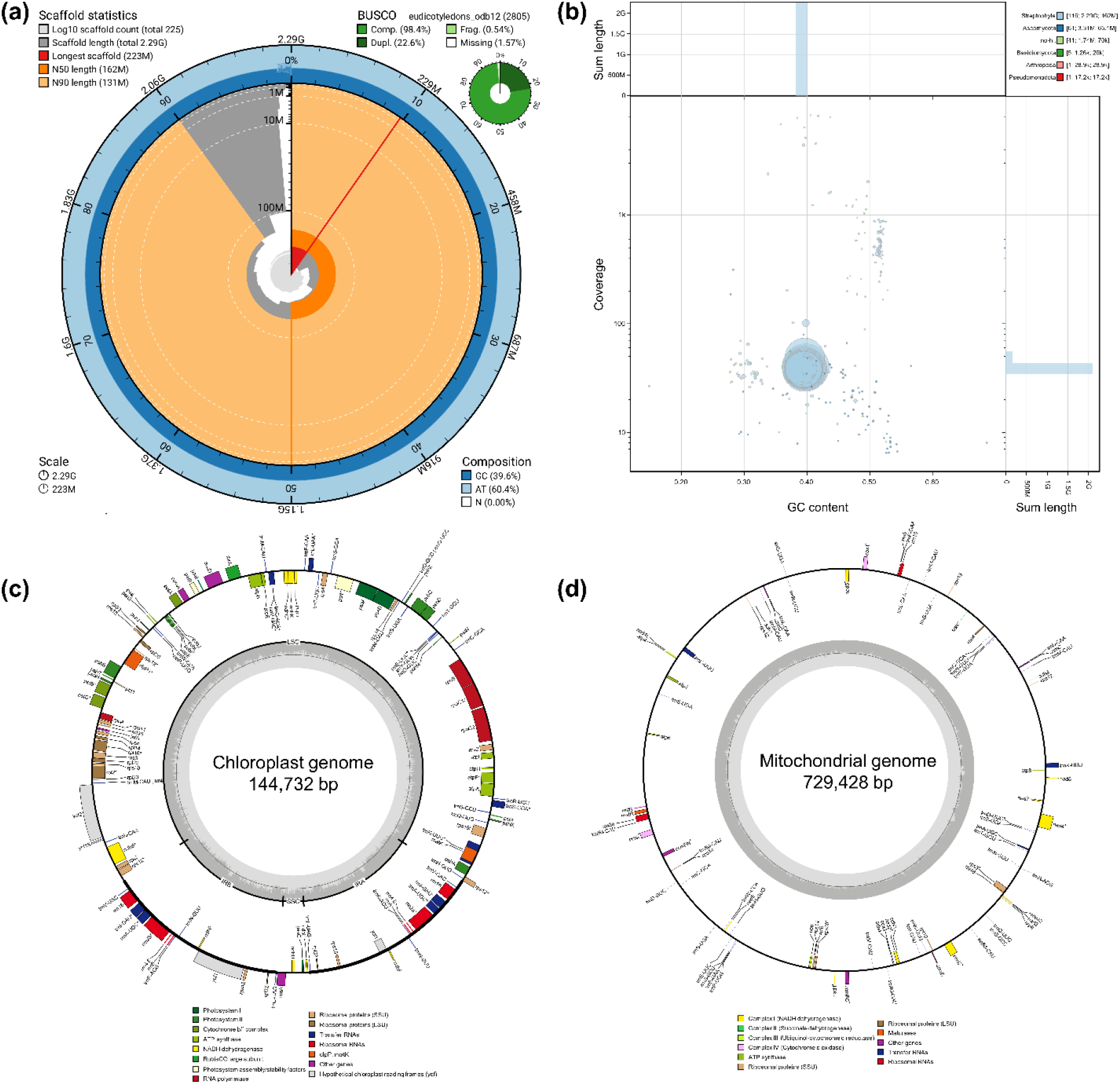
Nuclear genome statistics and organelle genome annotation of *Agalinis fasciculata*. (a) Scaffold statistics, BUSCO scores, and composition of the chromosome-level genome in a SnailPlot. (b) Sequencing coverage, sum length, and contamination across the genome in a BlobPlot. (c) Plastid genome structure and annotation, with genes colored by functional group. (d) Mitochondrial genome structure and annotation, with genes colored by functional group. For both (c) and (d), asterisks indicate genes containing introns.

### Organelle Genome Assembly

We assembled the plastid genome as a circular molecule of 144,732 bp (Fig. 1c). Genome size was consistent with that reported for autotrophic and hemiparasitic Orobanchaceae (Zhang et al. 2020). In holoparasitic Orobanchaceae, plastid genomes undergo extensive restructuring associated with the loss of photosynthesis, likely due to relaxed selection (Wicke et al. 2013), leading to substantial variation in genome size and gene loss. However, studies have indicated that the loss of tRNAs and *ndh* genes precedes the loss of photosynthesis in some hemiparasitic groups (Zhang et al. 2020). Consistent with this pattern, the plastid genome of *A. fasciculata* contains 109 genes, including 76 protein-coding genes, 29 tRNAs, and four rRNAs (Table S1), lacking some *ndh* genes present in other hemiparasitic Orobanchaceae species (Zhang et al. 2020). However, a more detailed screening is necessary to understand the plastid genome evolution and selective pressures in the *Agalinis* plastome.

The mitochondrial genome was assembled into a circular molecule of 729,428 bp (Fig. 1d), containing 61 genes, including 35 protein-coding genes, 23 tRNAs, and three rRNAs (Table S1). Fan et al. (2016) reported limited gene loss in the mitogenomes of parasitic plants. However, Cai et al. (2026) showed that non-core genes are often pseudogenized. For example, *sdh3* has multiple copies in *A. fasciculata*, all of which lack start or stop codons. Studies have also demonstrated that plant mitogenomes experience dynamic selective pressures (Feng and Wicke 2025) and exhibit an inverse relationship with plastid genome size in Orobanchaceae (Feng and Wicke 2025). The increasing availability of Orobanchaceae organelle genomes will enable a better understanding of how parasitism shapes organelle genome evolution, horizontal gene transfer, and interactions among the nuclear, plastid, and mitochondrial genomes, a promising area for future studies of parasitic lineages.

### Nuclear Genome Annotation and Tissue Specificity

BUSCO analysis (eudicotyledons_odb12) (Fig. 1a) indicated high genome completeness (98.4%), although with a high proportion of duplicated genes (22.6%). The genome of *A. fasciculata* is larger than that of *Castilleja foliosa* Hook. & Arn. (751.2 Mb), indicating that transposable element expansion contributes to its large genome size and high repetitive content (87.43%), particularly retroelements (Table S2), which accounted for 73.31% of the repeats. Both the genome size and repetitive content are consistent with the GenomeScope estimates (Fig. S1). Structural annotation predicted 34,133 protein-coding genes and 39,266 transcripts, which were reduced to 32,602 genes and 37,156 transcripts after functional annotation. Of these, 36,343 transcripts were assigned at least one functional annotation. Combined RNA data from flowers and flower buds exhibited the highest number of enriched genes, with most genes showing strong tissue specificity (Fig. S5A–B). When analyzed separately, flower buds and roots+haustoria had the highest number of enriched genes (Fig. S5C), although overall Tau was lower (Fig. S5D).

### Genetic Diversity, Runs of Homozygosity and Demographic History

Mean genome-wide heterozygosity in *A. fasciculata* was low (0.72 het./kb) and unevenly distributed across chromosomes, with higher levels observed on chromosomes 8 and 14 (Fig. 2A). Scans for runs of homozygosity (ROH), i.e., continuous regions of reduced heterozygosity, identified 467 long segments (≥1 Mb) of up to 5.96 Mb in length (Fig. 2B–C). Using a mutation rate of 2.12 × 10^−9^ per generation, demographic inference indicated that *A. fasciculata* reached its peak effective population size (*N*_*e*_) approximately 200,000 years ago, followed by a rapid decline until 20,000 years ago, when the population began a slow recovery (Fig. 2D). This recovery coincides with the Last Glacial Maximum, a period marked by the expansion of open vegetation habitats across southeastern North America (Hall and Valastro 1995), potentially facilitating population expansion of open vegetation species such as *Agalinis*. Although *A. fasciculata* is not currently considered a species of conservation concern, the low heterozygosity and *N*_*e*_ suggest high levels of inbreeding. While the reproductive biology of *A. fasciculata* remains unknown, *Agalinis* species are capable of self-fertilization, which may contribute to the low genetic diversity and high inbreeding levels.

**Figure 2.**
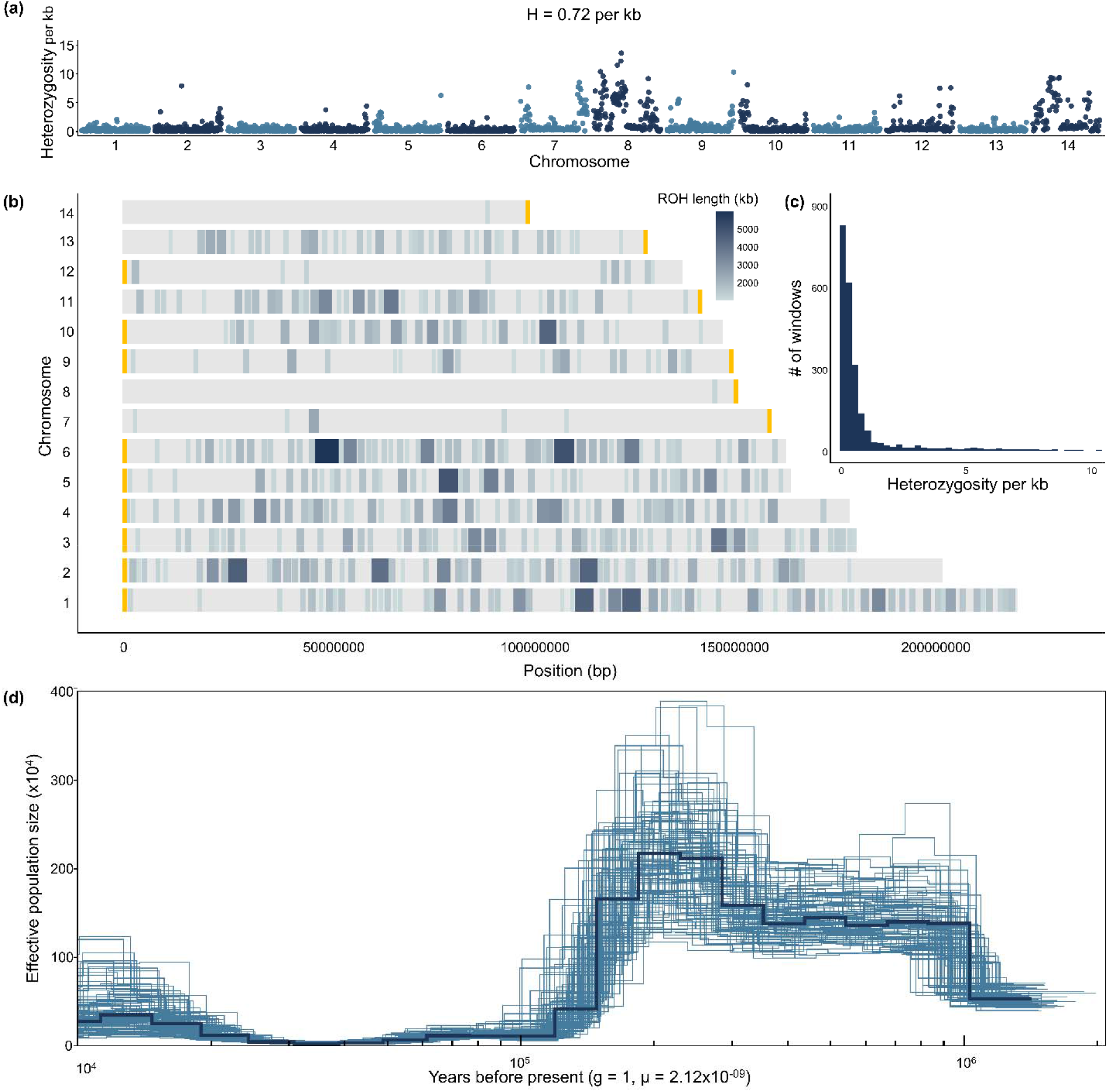
Genetic diversity and demographic history of *Agalinis fasciculata*. (a) Genome-wide heterozygosity estimated in non-overlapping 1-Mb windows across the 14 chromosomes. H indicates mean genome-wide heterozygosity. (b) Genomic location and length of runs of homozygosity (ROH) across the 14 chromosomes, with darker colors indicating longer ROH. Yellow bars at chromosome ends indicate telomeric repeats. (c) Frequency distribution and size of runs of homozygosity (ROH). (d) Historical changes in effective population size (N_e_) inferred for *A. fasciculata* using a mutation rate of 2.12 × 10□□per generation. Light blue lines represent 100 bootstrap replicates, whereas the dark blue line represents the estimate based on the whole genome.

## Materials and Methods

### DNA Extraction, PacBio HiFi and Omni-C Sequencing, and Quality Control

We collected several individuals of *A. fasciculata* at Cato Springs Prairie, University of Arkansas, Fayetteville (36°02′24.4″N, 94°10′59.9″W) in September of both 2024 and 2025, and snap-froze flowers, leaves, and roots+haustoria tissues in liquid nitrogen. A voucher specimen was deposited in the UARK Herbarium (accession no. UARK 095889). High-molecular-weight genomic DNA (gDNA) was extracted using a CTAB protocol and purified with a salt:chloroform wash. Library preparation was performed at the University of Oregon Genomics and Cell Characterization Core Facility (Eugene, OR), and sequencing was carried out on a PacBio Revio (Pacific Biosciences, Menlo Park, CA) using two Single Molecule, Real-Time (SMRT) cells. A different individual from the same population was used for Omni-C sequencing. Omni-C library preparation and sequencing were conducted at Dovetail Genomics (Scotts Valley, CA). Adapter sequences from HiFi reads were filtered using HiFiAdapterFilt (Sim et al. 2022). We used meryl (Rhie et al. 2020) to count 21-mers and used the output to estimate genome size, heterozygosity, and repeat content with GenomeScope 2.0 (Ranallo-Benavidez et al. 2020).

### Genome Assembly, Scaffolding, and Quality Assessment

We assembled the genome using hifiasm v0.19.5 (Cheng et al. 2021). Given the high level of homozygosity, downstream analyses were conducted on the primary assembly. We used purge_haplotigs v1.1.3 (Roach et al. 2018) to remove allelic variants from the primary assembly, with read depth cutoffs of -l 10, -m 60, and -h 200. Omni-C reads were mapped and filtered following the Dovetail Genomics reference guide.

We used YaHS v1.2.2 (Zhou et al. 2023) to scaffold our primary assembly. Plastid and mitochondrial scaffolds were removed by mapping the organelle genomes against the assembly using MUMmer v4.0.1 (Marçais et al. 2018), with a minimum identity of 95% and a minimum alignment length of 5,000 bp. To avoid removing nuclear mitochondrial DNA (NUMTs) and nuclear plastid DNA (NUPTs), we only filtered out scaffolds shorter than the corresponding organelle genome and with ≥75% coverage by that genome. We used Kraken v2.1.3 (Lu et al. 2022) to remove contaminant scaffolds, applying a confidence threshold of 0.3 with the PlusPFP database. We visualized contact maps using PretextSuite (https://github.com/wtsi-hpag/PretextView). We used tidk v0.2.65 (Brown et al. 2025) to identify telomeric repeat regions across scaffolds. As tidk detected telomeric repeats in the middle of the longest scaffold and karyotyping confirmed 14 chromosomes (Canne 1984), we manually split this scaffold to match the chromosome number. Assembly gaps were filled using YAGCloser (https://github.com/merlyescalona/yagcloser) with a flank size of 40 (-f 40) and a minimum of two reads (-mins 2) spanning each gap. We assessed genome quality and completeness using BUSCO v6.0.0 (Tegenfeldt et al. 2025) using the eudicotyledons_odb12 database. We assessed genome quality using BlobTools2 (Challis et al. 2020) and removed contaminant scaffolds that were kept by kraken2. We estimated ploidy level by analyzing heterozygous k-mer pairs with smudgeplot v0.4.0dev (Ranallo-Benavidez et al. 2020) and assessed genome quality with QUAST v5.3.0 (Gurevich et al. 2013). We used Oatk v1.0 (Zhou et al. 2025) to assemble the organelle genomes. The plastid genome was annotated using GeSeq (Tillich et al. 2017), and the mitogenome was annotated using PMGA (Li et al. 2025). Organelle genomes were visualized with OGDRAW v1.3.1 (Greiner et al. 2019).

### RNA Sequencing, Genome Annotation, and Tissue Specificity

We generated a de novo, species-specific repetitive sequence and transposable element (TE) library using RepeatModeler2 v.2.0.7 (Flynn et al. 2020). Sequences labeled as “unknown” were classified following Nachtigall et al. (2025). Briefly, we used DeepTE v1.0 (Yan et al. 2020) with the plant-specific model to classify those repetitive sequences and removed false positives using TERL v1.0 (Da Cruz et al. 2021). We used this dataset to soft-mask repetitive regions in the genome with RepeatMasker v4.2.1 (Smit et al. 2013; https://www.repeatmasker.org). We soft-masked NUMTs and NUPTs by using the assembled organelle genomes as queries in a BLASTn search against the nuclear genome.

Flower, flower bud, leaves, and root+haustoria tissues were sent to Novogene (Sacramento, CA) for RNA extraction, library preparation, and sequencing on an Illumina NovaSeq X Plus. We used fastp v1.0.1 (Chen et al. 2018) to process RNA-seq reads, followed by a correction step using Rcorrector v1.0.7 (Song and Florea 2015). We mapped the RNA-seq reads to the assembly using HISAT2 v2.2.1 (Kim et al. 2019) and used its output for structural genome annotation with BRAKER3 v3.0.8 (Gabriel et al. 2024). We also used the Viridiplantae protein dataset from OrthoDB v12 (Tegenfeldt et al. 2025) and the proteomes of nine Orobanchaceae species (Table S3) as protein evidence.

We used eggNOG-mapper v2.1.12 (Cantalapiedra et al. 2021; http://eggnog-mapper.embl.de/) and InterProScan v5.59-91.0+galaxy3 (Jones et al. 2014) via Galaxy Europe (Afgan et al. 2018; https://usegalaxy.eu/) for functional annotation. The outputs were then used as input for Funannotate v1.8.17 (Palmer and Stajich 2020). Functional annotation was supplemented with BLASTp searches for genes labeled as hypothetical proteins. We used three protein datasets (retrieved on 15 Oct 2025) hierarchically: 1) a reviewed and curated SwissProt protein dataset of eudicotyledons (28,390 sequences); 2) an unreviewed TrEMBL dataset of lamiid proteins (2,376,208 sequences); and 3) a curated SwissProt dataset across all taxa (573,661 sequences). We used STAR v.2.7.11b (Dobin et al., 2013) to align the tissue RNA-seq reads to the novel genome. We compared expression across the tissues to understand transcript distribution and tissue specificity using the Tau (Yanai et al. 2005)

### Genetic Diversity and Population Size History

As in Gonçalves et al. (2025), we evaluated genome-wide heterozygosity and ROH as proxies for genetic diversity in *A. fasciculata*. Filtered HiFi reads were mapped to the assembled genome using minimap2 (Li 2018). The resulting BAM file was sorted with SAMtools (Danecek et al. 2021), and a VCF file containing variants and invariants was created using BCFtools v1.22 (Danecek et al. 2021) with minimum base and mapping quality thresholds of 30 (-Q 30 and -q 30). We used pixy v2.0.0.beta14 (Korunes and Samuk 2021) to estimate nucleotide diversity across non-overlapping 1 Mb sliding windows and PLINK v1.9.0-b.8 (Chang et al. 2015) to estimate ROH.

We used the Pairwise Sequentially Markovian Coalescent model (PSMC; https://github.com/lh3/psmc) to infer historical changes in *N*_*e*_ of *A. fasciculata*. PSMC was run with 100 bootstrap replicates using the following parameters: -N25 -t15 -r5 -p “1+1+1+1+25*2+4+6”. A mutation rate was estimated by calculating the number of 1:1 genomic matches between *A. fasciculata* and *C. foliosa* (Bürger et al. 2024). The masked genomes were aligned with nucmer from MUMmer and the output was filtered using delta-filter via the Galaxy Europe platform. The total number of SNPs was divided by the number of 1:1 matches to estimate overall genomic divergence, which was converted into a mutation rate per generation using the formula described in Luo et al. (2023), assuming a mean generational time of 1 year and a divergence time between *Agalinis* and *Castilleja* of 32.93 mya (Fonseca 2021). These analyses were restricted to the 14 chromosomes.

## Acknowledgments

Computational analyses were supported by the Arkansas High Performance Computing Center, which is funded through multiple National Science Foundation grants and the Arkansas Economic Development Commission. We also thank Jennifer Ogle, M.S., for assistance with sampling and species identification, Sarah Jacobs, Ph.D., for advice on DNA extraction, and Leonardo Tresoldi Gonçalves, Ph.D., for support with sampling and bioinformatics analyses.

## Funding

This research was supported by the University of Arkansas start-up research funds to M. L.

## Data availability

Scripts with all the parameters and files used for the analyses are available on GitHub https://github.com/pedrohpezzi/Genome_Agalinis_fasciculata and https://doi.org/10.5281/zenodo.18393392. Raw reads and genome assembly have been deposited in GenBank under the BioProject accession number PRJNA1345713.

## CRediT Author Contribution

PHP: Conceptualization, Data curation, Formal Analysis, Investigation, Methodology, Software, Validation, Visualization, Writing – original draft, Writing – review & editing; ML: Conceptualization, Funding acquisition, Project administration, Resources, Supervision, Writing – review & editing.

## Conflict of Interest

The authors declare that they have no conflicts of interest.

---

**Figure S1.**
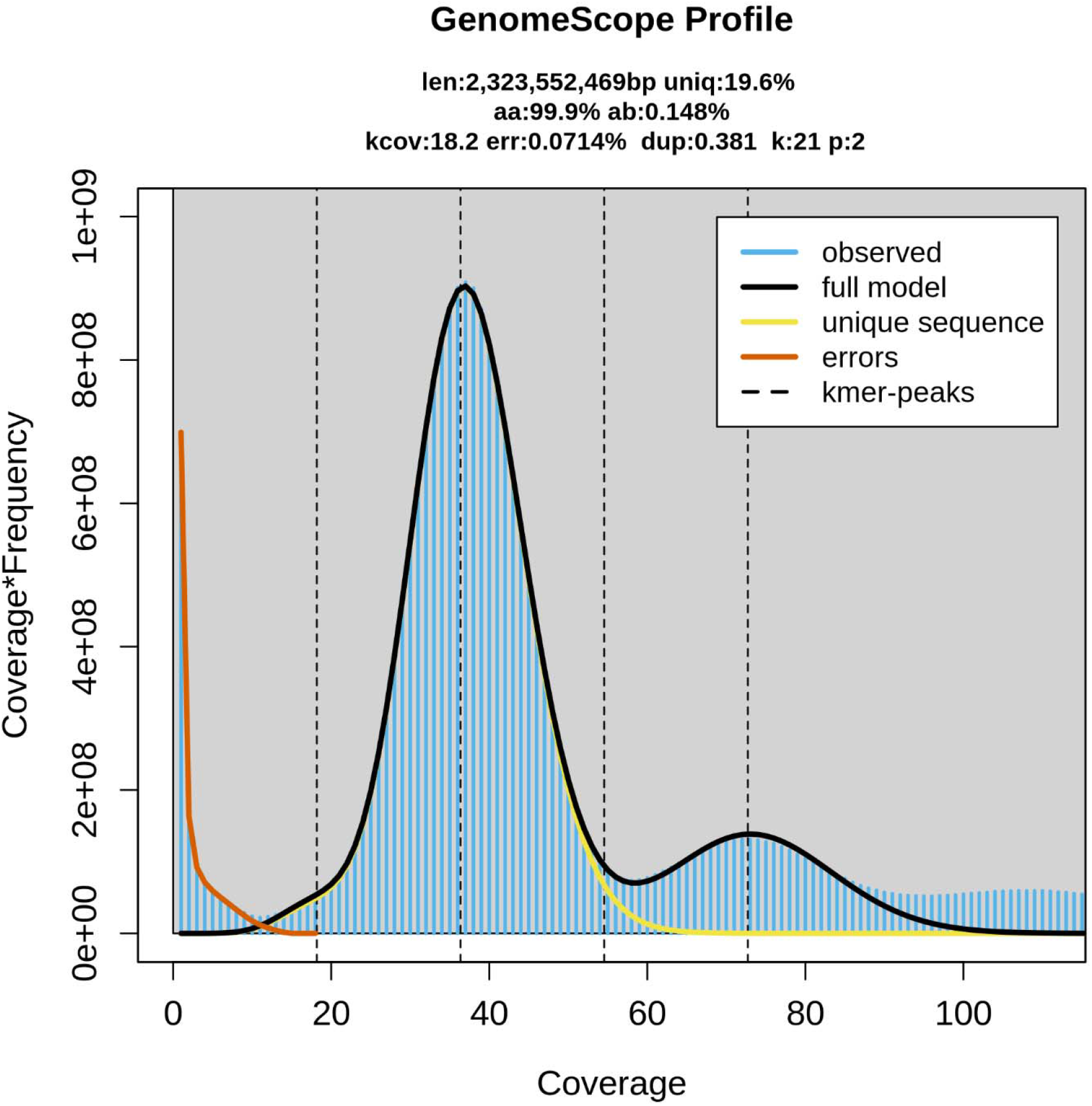
GenomeScope profile of *Agalinis fasciculata* based on 21-mers. The k-mer frequency distribution shows two peaks, typical of diploid genomes. GenomeScope estimates a genome size of ∼2.3 Gb with low heterozygosity (0.15%).

**Figure S2.**
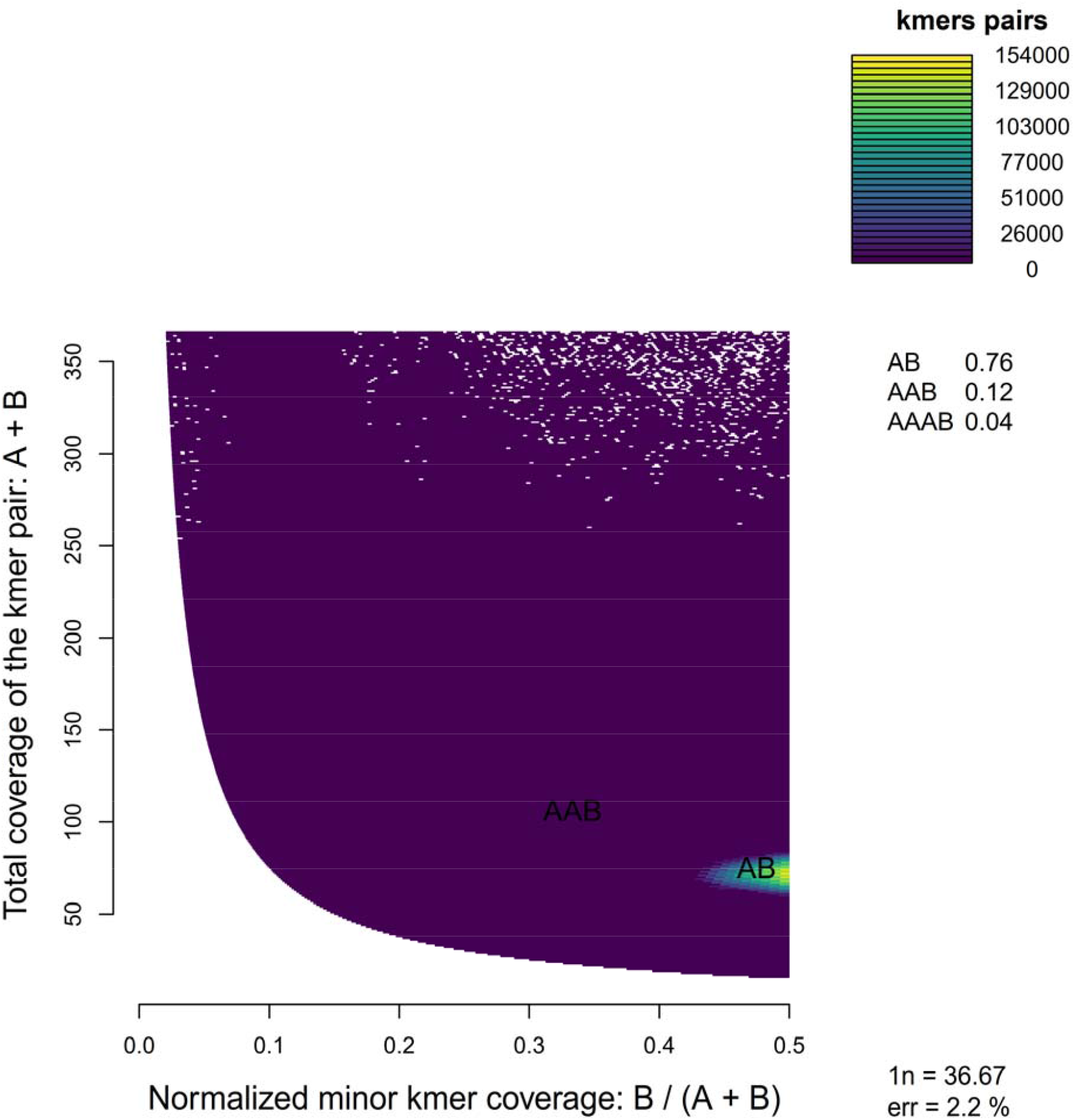
Smudgeplot estimation of the ploidy level of *Agalinis fasciculata* based on 21-mers. The dominant AB smudge (0.76) is consistent with a diploid genome.

**Figure S3.**
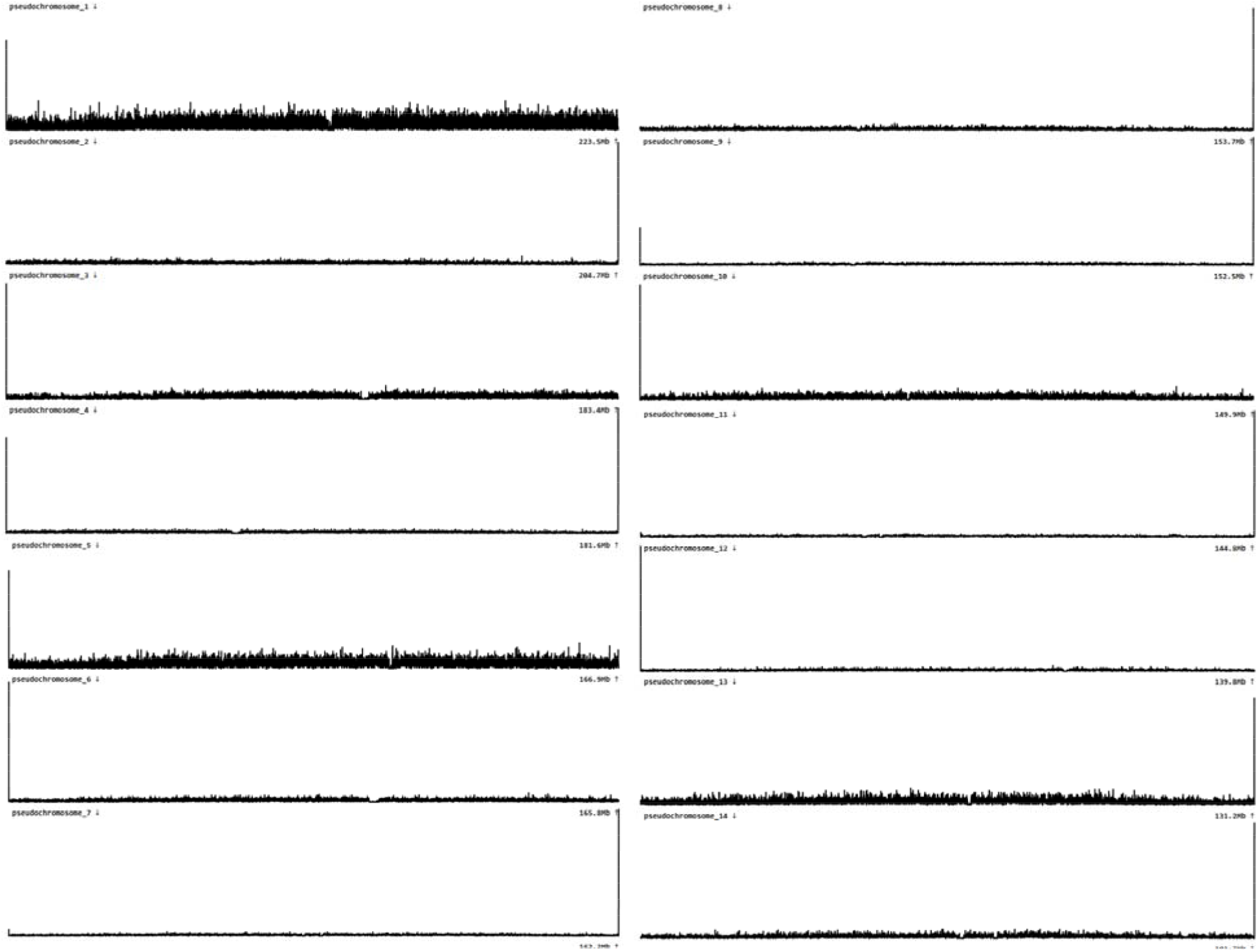
Telomeric repeats identified in the 14 chromosomes of *Agalinis fasciculata* using tidk. All chromosomes show at least one telomeric peak at one or both chromosome ends. Telomeric repeats are indicated by peaks in the histograms.

**Figure S4.**
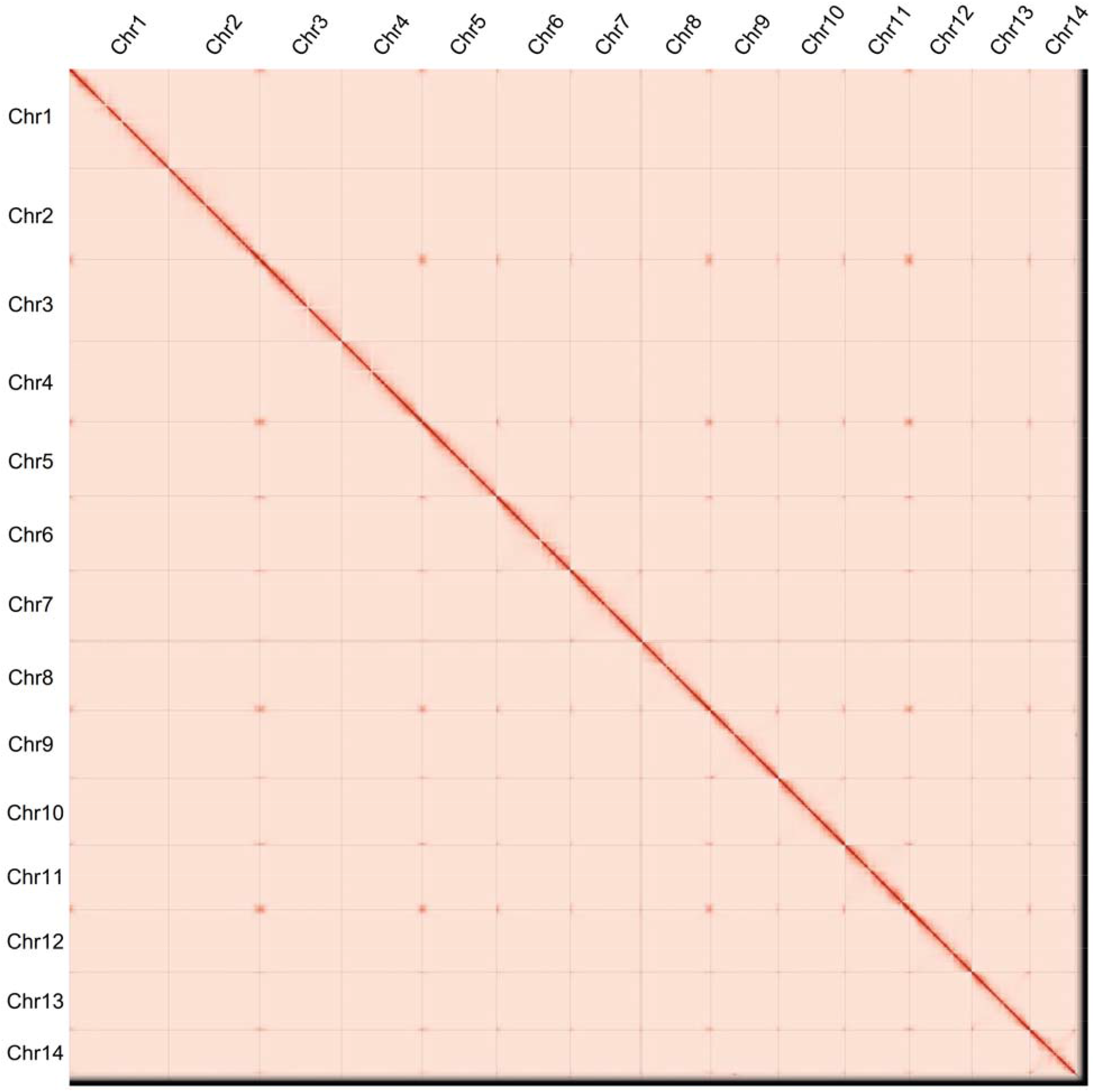
Omni-C contact map of the chromosome-level assembly of *Agalinis fasciculata*. The contact map shows strong intrachromosomal interactions (red diagonal) across the 14 chromosomes.

**Figure S5.**
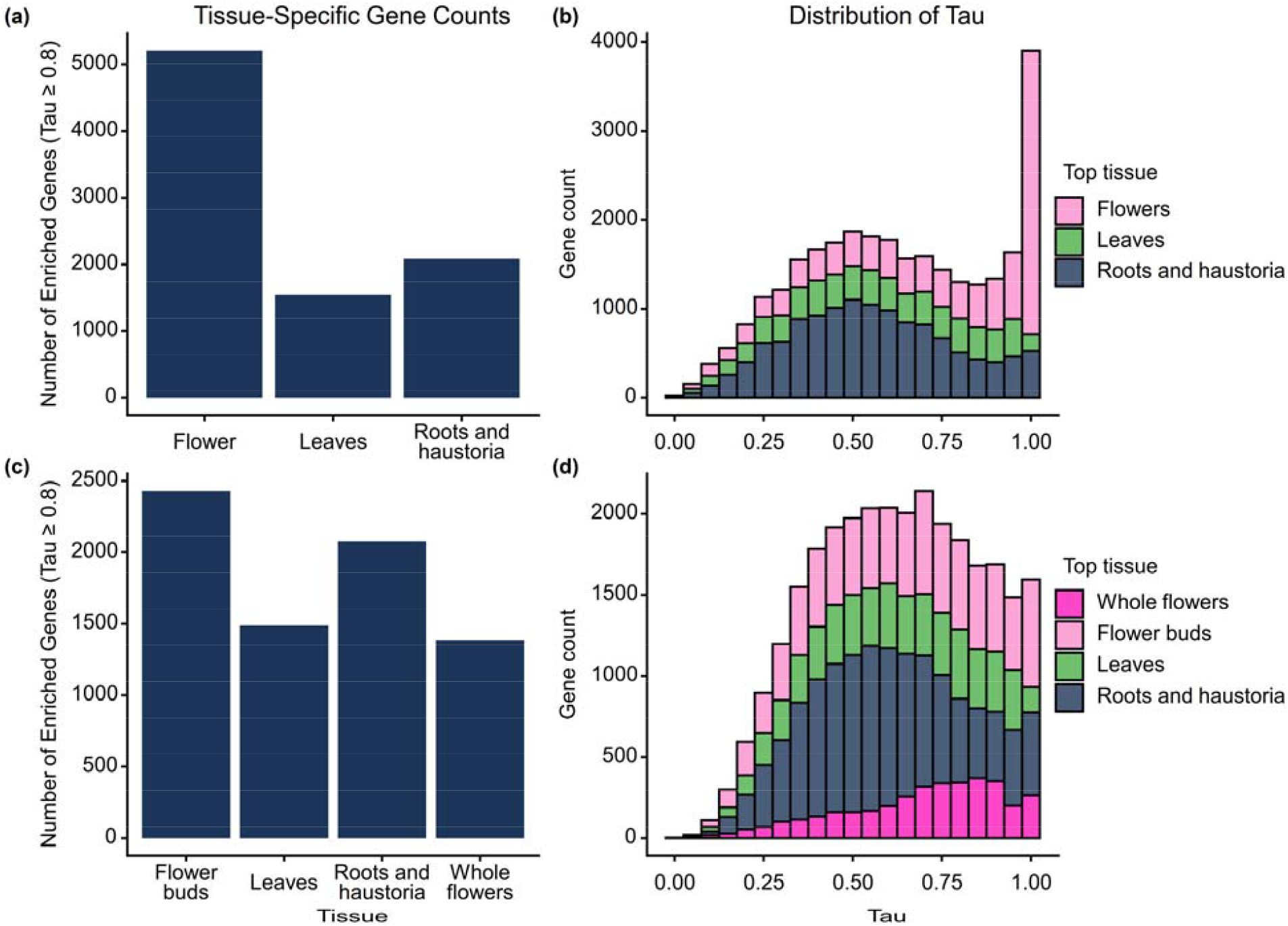
Tissue-specific gene expression in *Agalinis fasciculata*. Tissue specificity (Tau) of gene expression based on RNA-seq data, shown for flower and flower bud tissues combined (a– b) and for the four tissues analyzed separately (c–d). For (a) and (c), genes with Tau ≥ 0.8 were considered enriched.

**Table S1.**
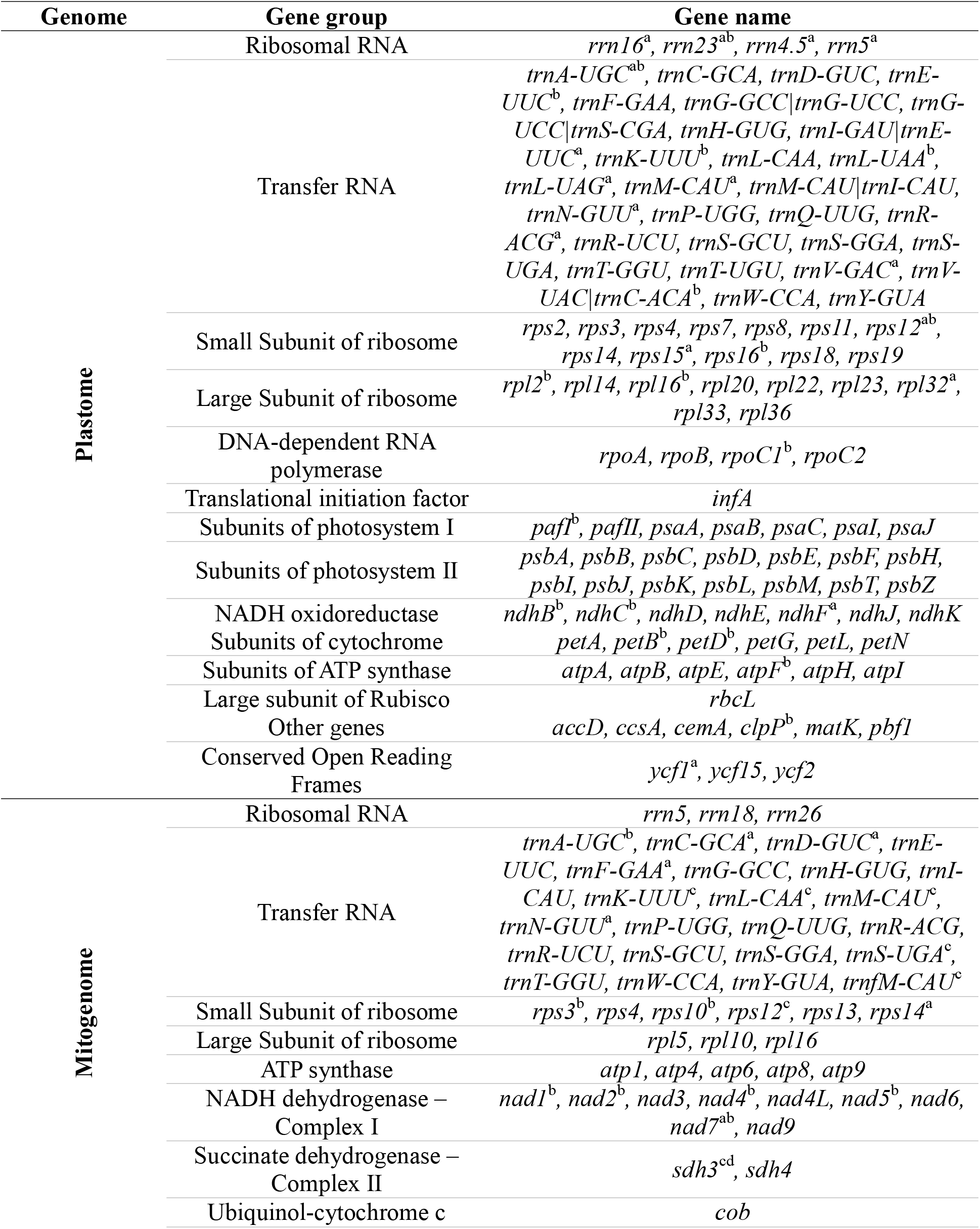
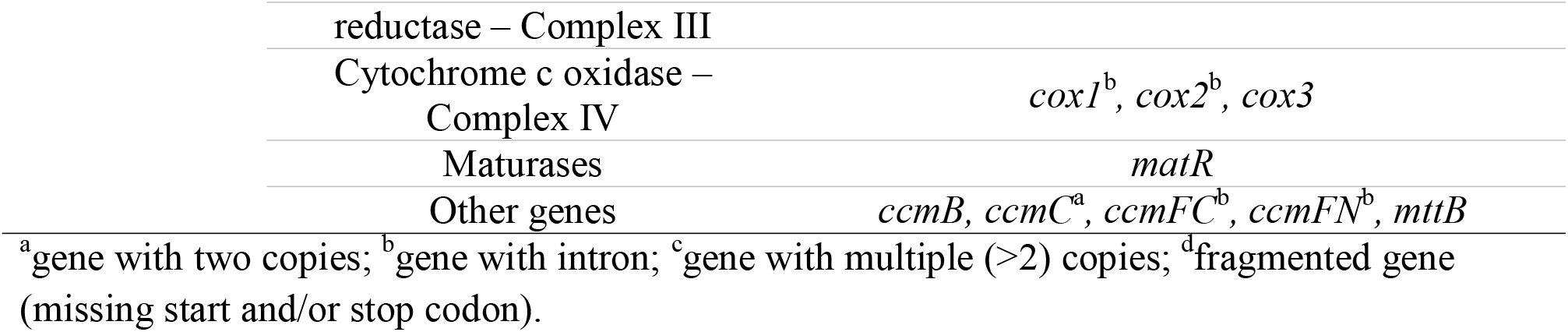
Genes present in the plastid (plastome) and mitochondrial (mitogenome) genomes of *Agalinis fasciculata*, grouped by functional category.

**Table S2.** Number, total length, and genomic proportion of repetitive elements in the *Agalinis fasciculata* genome, as classified by RepeatMasker.

| Class | Order | Superfamily | Number of Elements | Length occupied (bp) | Genome % |
| --- | --- | --- | --- | --- | --- |
| **Retroelements** | - | - | 977,747 | 1,676,710,586 | 73.31 |
|  | LINEs | - | 38,114 | 22,465,310 | 0.98 |
|  | LINEs | RTE/Bov-B | 9,793 | 2,646,803 | 0.12 |
|  | LINEs | L1/CIN4 | 28,321 | 19,818,507 | 0.87 |
|  | LTR elements | - | 939,633 | 1,654,245,276 | 72.33 |
|  | LTR elements | Ty1/Copia | 208,840 | 546,489,247 | 23.89 |
|  | LTR elements | Gypsy/DIRS1 | 711,227 | 1,089,729,195 | 47.65 |
| **DNA transposons** | - | - | 379,953 | 213,114,117 | 9.32 |
|  | hobo-Activator | - | 170,622 | 75,436,725 | 3.30 |
|  | Tc1-IS630-Pogo | - | 49,030 | 16,351,179 | 0.71 |
|  | MULE-MuDR | - | 24,384 | 17,844,775 | 0.78 |
|  | Tourist/Harbinger | - | 47,907 | 22,297,828 | 0.97 |
| **Rolling-circles** | - | - | 1,771 | 989,873 | 0.04 |
| **Unclassified** | - | - | 161,288 | 50,508,830 | 2.21 |

**Table S3.** Proteomes from Orobanchaceae species used for structural annotation, with corresponding GenBank accession numbers.

| **Species** | **Accession number** |
| --- | --- |
| *Castilleja foliolosa* | GCA_046119335.1 |
| *Orobanche gracilis* | GCA_050575165.1 |
| *Orobanche hederæ* | GCA_050575145.1 |
| *Orobanche minor* | GCA_050575125.1 |
| *Phtheirospermum japonicum* | GCA_014905375.1 |
| *Rehmannia glutinosa* | GCA_016081115.2 |
| *Striga asiatica* | GCA_008636005.1 |
| *Striga hermonthica* | GCA_902706635.1 |

## Notes

https://github.com/pedrohpezzi/Genome_Agalinis_fasciculata

https://doi.org/10.5281/zenodo.18393392

### Competing Interest Statement

The authors have declared no competing interest.

